# Unbiased Complete Estimation of Chloroplast Number in Plant Cells Using Deep Learning Methods

**DOI:** 10.1101/2023.12.17.572064

**Authors:** Qun Su, Le Liu, Zhengsheng Hu, Tao Wang, Huaying Wang, Qiuqi Guo, Xinyi Liao, Zhao Dong, Shaokai Yang, Ningjing Liu, Qiong Zhao

## Abstract

Chloroplasts are essential organelles in plants that are involved in plant development and photosynthesis. Accurate quantification of chloroplast numbers is important for understanding the status and type of plant cells, as well as assessing photosynthetic potential and efficiency. Traditional methods of counting chloroplasts using microscopy are time-consuming and face challenges such as the possibility of missing out-of-focus samples or double counting when adjusting the focal position. Here, we developed an innovative approach called Detecting- and-Counting-chloroplasts (D&Cchl) for automated detection and counting of chloroplasts. This approach utilizes a deep-learning-based object detection algorithm called You-Only-Look-Once (YOLO), along with the Intersection Over Union (IOU) strategy. The application of D&Cchl has shown excellent performance in accurately identifying and quantifying chloroplasts. This holds true when applied to both a single image and a three-dimensional (3D) structure composed of a series of images. Furthermore, by integrating Cellpose, a cell-segmentation tool, we were able to successfully perform single-cell 3D chloroplast counting. Compared to manual counting methods, this approach improved the accuracy of detection and counting to over 95%. Together, our work not only provides an efficient and reliable tool for accurately analyzing the status of chloroplasts, enhancing our understanding of plant photosynthetic cells and growth characteristics, but also makes a significant contribution to the convergence of botany and deep learning.

**One-sentence summary:** This deep learning-based approach enables the accurate complete detection and counting of chloroplasts in 3D single cells using microscopic image stacks, and showcases a successful example of utilizing deep learning methods to analyze subcellular spatial information in plant cells.

The authors responsible for distribution of materials integral to the findings presented in this article in accordance with the policy described in the Instructions for Authors (https://academic.oup.com/plcell/) is: Zhao Dong, Shaokai Yang, Ningjing Liu, and Qiong Zhao.

## Introduction

Chloroplasts convert light energy to chemical energy through photosynthesis, provide oxygen, and act as a cornerstone for the world (Whatley, 1975). The chloroplasts were originated approximately 1.5 billion years ago, when a prokaryotic photosynthetic bacterium ancestor was engulfed by a eukaryotic cell in a singular endosymbiotic event (Archibald, 2009; Keeling, 2013). Studies on chloroplasts mainly focus on its material composition, genome, diversity and evolution, structures, as well as function and adaption (Schubert et al., 2002; van Wijk et al., 2007; Demartini et al., 2011; Gros and Jouhet, 2018; Wakasugi et al., 1997; Gray et al., 1999; Kirchhoff, 2019; Daniell et al., 2016; Cavalier-Smith, 2002; Song et al., 2021; Ouyang et al., 2020). Typically, chloroplasts are elliptical or disc-shaped structures, with pigments accumulated inside, resulting in a green color under microscopic observation (Murata, 1969; Kume, 2017). Studies showed that the number of chloroplasts might be associated with cell status or function, suggesting a correlation between chloroplast count and plant cell type (Pyke and Leech, 1994; Ii and Webber, 2005). For instance, the reduction in chloroplast number affects both the composition and structure of the photosynthetic apparatus, suggesting that the ability for photosynthesis relies on proper chloroplast division and development (Li and Webber, 2005). Understanding the mechanism of chloroplast number coordination in a specific cell type is a fundamental inquiry. Stomatal guard cells in the plant shoot epidermis typically possess several to tens of chloroplasts per cell (Macfarlane, 1898; Sakisaka, 1929; Mochizuki and Sueoka, 1955; Fujiwara et al., 2019). So far, the number of chloroplasts at the stomatal level has commonly been utilized as a convenient indicator for identifying hybrid species or estimating the ploidy level of a particular plant tissue (Fujiwara et al., 2019; Watts et al., 2023). Ukwueze et al. investigated the ploidy of banana germplasm through chloroplast counting methods, while Chepkoech et al. (2019) observed an increase in chloroplast numbers in tetraploid potatoes compared to the diploid plant (Ukwueze et al., 2022; Chepkoech et al., 2019). Moreover, Pyke and his colleagues found that the average chloroplast number per cell in the first leaves differs among different genetic backgrounds of *Arabidopsis* (Pyke and Leech, 1994; Pyke et al., 1994). In the Landsberg erecta (Ler) ecotype, the number is 121, while in the Wassilewskija (Ws) ecotype, it is 83 (Pyke and Leech, 1994; Pyke et al., 1994). The above studies suggest the number of chloroplasts can act as a classification marker.

Till now, the most commonly used counting methods in the aforementioned studies have been manual counting using images captured via light-microscope, which is time-consuming and prone to misidentification. Molecular staining applied to organelles counting (including chloroplast) associated with flow cytometry may sound high-flux, but it is restricted to the isolation process and cannot accurately determine the number per cell (Mattiasson, 2004; Cole, 2016). Chloroplast counting towards *Spruce needle* leaf showed that detection on the three-dimensional (3D) structure of mesophyll cells, obtained by continuous optical cross-sections via confocal laser-scanning microscopy, provides a wealth of information beyond traditional two-dimensional (2D) images (Kubínová et al., 2014). The authors found that approximately 90% of chloroplasts were lost in the 2D images compared to the 3D structure, when the cells were thick (Kubínová et al., 2014). Yet, this method relies on complex data collection, which can be influenced by subjectivity and technical constraints. Semi-automatic cell counting methods were employed, which relied on manual-thresholding segmentation and automated measurement using expert software, such as ImageJ (Arena et al., 2017). However, overlapped cells may not be accurately distinguished and counted.

As we enter the age of intelligence, deep-learning-based tools in life sciences have become widely utilized. Deep learning, in essence, mimics the learning process of the human brain and has its roots in the study of artificial neural networks. However, due to technological limitations, its progression was stymied for a while. It wasn’t until 2012 that the field saw a game-changing moment: Krizhevsky, Sutskever, and Hinton, with their AlexNet, claimed victory in the ImageNet competition, surpassing all other approaches (Krizhevsky, Sutskever and Hinton, 2012). They harnessed the power of GPUs for accelerated computation, thereby demonstrating the remarkable potential and efficacy of deep learning in handling large-scale datasets. This breakthrough provided invaluable inspiration to subsequent researchers, catalyzing the widespread adoption of deep learning across a myriad of application domains. Particularly in cell biology, deep learning has unveiled remarkable potential in live cell imaging experiments. Van Valen and his colleagues harnessed the power of deep convolutional neural networks to segment and analyze individual cells in microscopic images, notably achieving high precision in the cytoplasmic segmentation of mammalian cells (Van Valen et al., 2016). In the realm of chemistry, deep learning has paved new pathways for innovative compound design. Utilizing variational autoencoder techniques, scientists have successfully transformed the discrete representations of molecules into continuous multi-dimensional spaces, allowing for the automated generation of innovative chemical structures. This provides a powerful tool for the efficient exploration and optimization of vast chemical compound spaces. (Gómez-Bombarelli et al., 2018). Furthermore, deep learning has been adeptly employed in predicting the sequence specificity of DNA and RNA binding proteins. Leveraging a technique known as DeepBind, researchers are equipped to autonomously discern novel sequence patterns and compute predictive binding scores. Impressively, its performance surpasses that of other existing methods across various experimental datasets (Alipanahi et al., 2015).

By utilizing convolutional neural networks (CNNs) to estimate the spatial density map of cells, Xie et al. successfully achieved automated cell counting and detection in microscopy images (Xie et al, 2018). Transfer learning and a CNN were combined to analyze over 47,000 confocal fluorescence images from Arabidopsis, resulting in the development of a deep-learning framework called DeeplearnMOR (Deep Learning of the Morphology of Organelles). This tool enables rapid classification of image categories and accurate identification of abnormalities in organelle morphology (Li et al., 2021). An ImageJ plugin that enables non-machine-learning experts to analyze their data with U-Net was provided to solve for frequently occurring quantification tasks such as cell detection and shape measurements in biomedical image data (See http://sites.imagej.net/Falk/plugins/; Falk et al., 2019). You Only Look Once (YOLO) is a popular object detection algorithm that revolutionized the field of computer vision (Redmon and Farhadi, 2016). In YOLO, a single neural network is applied to the entire image, dividing it into a grid. This enables YOLO to make real-time predictions with impressive speed, as it only needs a single pass through the network to detect objects in an image, making it an excellent choice for cell detecting and counting (Redmon et al., 2016a; Redmon and Farhadi, 2016b; Alam and Islam,2019; Aldughayfiq et al., 2023). However, to the best of our knowledge, there have been no applications of this method to quantify subcellular structures such as chloroplasts or mitochondria using 3D images obtained by optical microscopes from living cells.

Here, we developed a YOLOv7-based deep learning tool called Decting&Counting-chloroplast (D&Cchl), integrated with the concept of Intersection Over Union (IOU), for accurate chloroplast counting in light microscope-captured images. Moreover, we employed the segmentation tool Cellpose to enable counting within individual 3D living cells (Pachitariu and Stringer, 2022). Overall, we achieved the precise complete 3D cell chloroplast counting.

## Results

### Work frame of the D&Cchl Model for plant cell 3D living cell chloroplasts detection and counting

To train our model, bright-field images of three bryophyte (*Sphagnum squarrosum* [*S. squ*], *Physcomitrium patens* [*P. pat*], and *Ricciocarpos natans* [*R. nat*]) leaf were obtained under a light microscope (Supplemental Fig. S1). As ideal plant materials, these plant leaves are single-cell-layered with regular cell distribution (Supplemental Fig. S1). Totally, 310 pictures (45 of *S. squ*, 56 of *P. pat*, 209 of *R. nat*) were obtained for chloroplast dataset collection. The software **LabelImg** (See https://github.com/tzutalin/labelImg; Tzutalin, 2015) was used for data pre-treatment to create well-manual-labelled dataset for subsequent model training (Tzutalin, 2015). Then labeled images were fed to the YOLOv7 framework loaded with Yolov7.pt model (Wang et al., 2023). After 500 round 2.662 h training on an NVIDIA GeForce RTX 3060 with 12GB of VRAM, we achieved a new model named D&Cchl (Fig. 1A).

**Figure 1.**
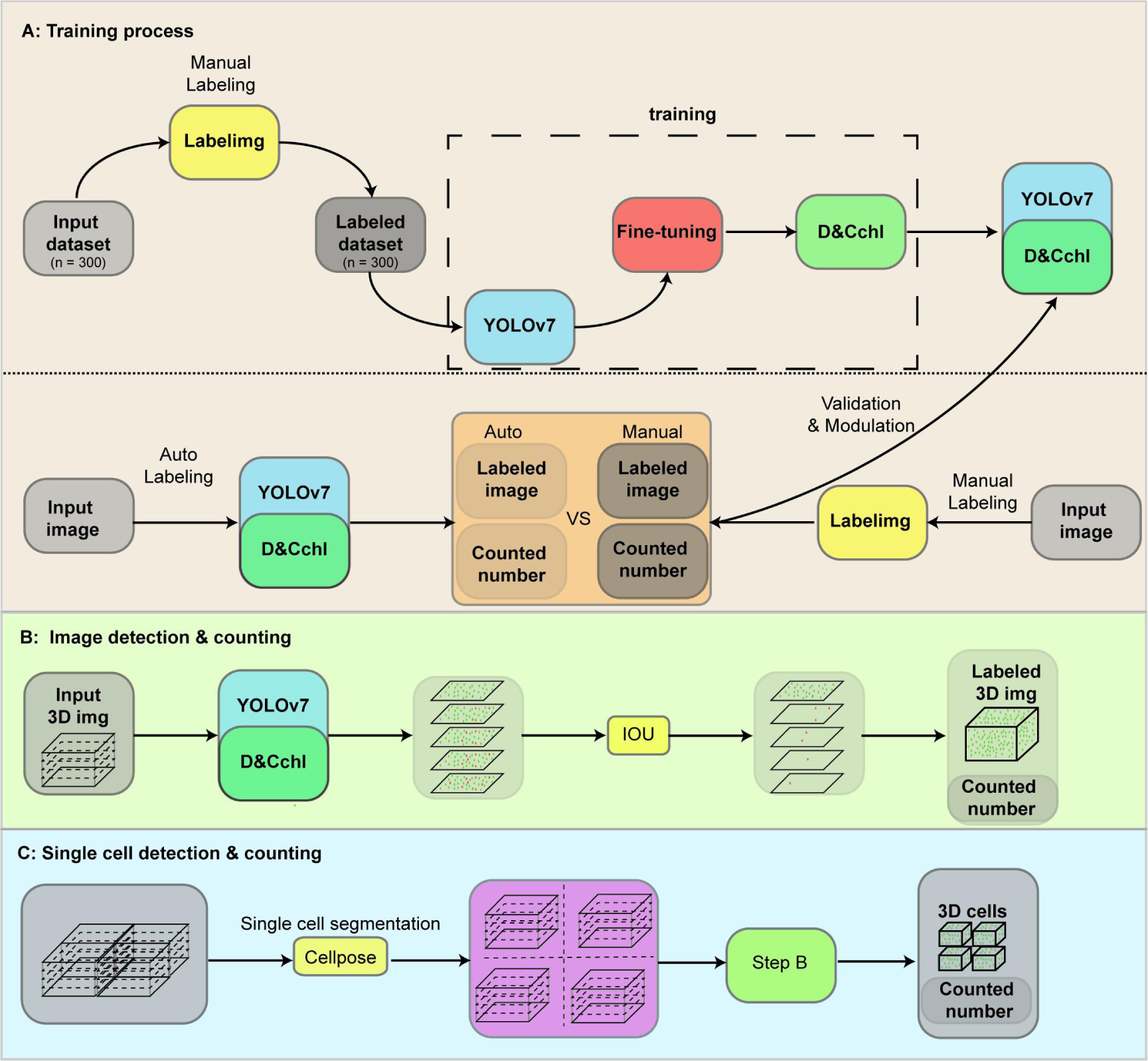
Workflow of the *YOLOv7-D&Cchl* for whole-cell chloroplast detection and counting. **A)** Detailed steps for the ***YOLOv7-D&Cchl*** training process. Microscopic images of plant cells were collected and chloroplasts were first labeled manually using ***Labelimg***. Then, the labeled dataset was further divided into training and validation sets to train ***YOLOv7*** for the chloroplasts auto-detection and counting function of the ***YOLOv7-D&Cchl*** model. Furthermore, the manually annotated data was compared with the model’s predictions. Based on these comparisons, adjustments were made to the model’s parameters to improve its accuracy. **B)** Strategy and procedure for detecting and counting chloroplasts in a 3D volume composed of a stack of multiple 2D layers. Accurate counting of chloroplasts in a 3D volume was achieved by using five continuous images spanning an entire cell from different focal planes as input. To prevent duplicate counting, the ***Intersection over Union*** *(**IOU**)* commutating mode was developed and integrated. **C)** Process for detecting and counting chloroplasts in individual 3D volume cells. Cell segmentation was performed by feeding continuous images spanning an entire cell from different focal planes into the pre-trained ***Cellpose*** model. The resulting segmented images were then inputted into the trained ***D&Cchl*** model for chloroplast detection. After the ***IOU*** was computed, the detection and counting of chloroplasts in the single cell 3D volume were achieved. Blocks colored in gray represents the dataset, yellow represents software, blue represents the neural network framework, and dark green represents the models generated from the training process. Text in ***italic bold*** represents program/model.

As widely recognized, detecting chloroplasts within 3D objects significantly enhances the accuracy of chloroplast counting per cell. In this study, we developed an IOU module to achieve accurate counting within a 3D volume. The module helps reduce repeated counting over a series of 2D images. We set a threshold value of 0.3 for IOU to measure the overlap between the proposed boxes and the reference boxes. The value of IOU ranges from 0 to 1, indicating the degree of overlap between the detected box and the true label, with 0 indicating no overlap and 1 indicating complete overlap (Fig. 1B). We selected five images from different focal planes as input for three-dimensional (3D) image detection and counting. It’s worth noting that during the data collection process, there were almost no lateral shifts in the images during the focusing process. With the assistance of the D&Cch model, we conducted chloroplast detection and generated detection results for each image. Starting from the first image, all detected chloroplasts were considered as a benchmark. For each subsequent image, the detected chloroplasts were compared with those in the benchmark layer based on their IOU. If the IOU exceeded a preset threshold, it was classified as an existing chloroplast from the benchmark layer, avoiding recounting. Otherwise, it was identified as a new chloroplast and added to the benchmark. The threshold primarily depended on the experiment. If there was minimal lateral image shift during the focusing process, a smaller threshold should be used. Coupled with our proposed IOU method, this approach not only improved potential errors stemming from two-dimensional (2D) image detection but also significantly sped up the processing time compared to manual counting.

Finally, to further enhance the precision and practicality of chloroplast counting within single 3D cell, we employed the Cellpose model to segment the images on a single-cell basis (Fig. 1C; Supplemental Fig. S2). This allowed us to achieve 3D detection on individual cells and perform accurate counting of chloroplasts within each cell. By using this advanced technology, we were able to improve the accuracy of our chloroplast counting method significantly. The trained Cellpose model (cyto2) were modified with 11 *R. nat* images to create the suitable model Cyto2Pro, enabling a more detailed and comprehensive analysis of chloroplast distribution within the cells. This breakthrough in chloroplast counting methodology opens up new possibilities for studying the role of chloroplasts in various cellular processes.

### The YOLOv7-D&Cchl demonstrated good performance in counting and detecting chloroplasts using 2D images

To thoroughly assess the performance of YOLOv7-D&Cchl in chloroplast detection, we employed the comprehensive evaluation metric, mAP, which combines precision and recall (Fig. 2A; Supplemental Fig. S3). In the validation dataset, when the confidence level was set to 0.5, the model achieved an average precision of 0.877. Furthermore, the F1 curve peaked at 0.84, further demonstrating the model’s excellent balance between precision and recall (Supplemental Fig. S4; Supplemental Fig. S5). Additionally, we manually labeled 9 images (3 images from each different plant) with complete chloroplast as a test set to evaluate the efficiency of YOLOv7-D&Cchl in detecting chloroplast (Fig. 1B; Fig. 2B). The model successfully detected 88 chloroplasts in *S. squ* (missing 3), 160 chloroplasts in *P. pat* (missing 10), 194 chloroplast in *R.nat* (missing 20), and 475 chloroplast in total images (missing 33), while falsely detected 0 chloroplasts in *S. squ*, 0 chloroplasts in *P. pat*, 0 chloroplast in *R.nat*, and 0 chloroplast in total (Fig. 2C). We also obtained the final counting results and found the precision rates are 97.10%, 96.77%, and 92.72% respectively (Fig. 2D; Supplemental Fig. S6; Supplemental Fig. S7). In all, the D&Cchl model has showed a good performance on chloroplast detecting and counting.

**Figure 2.**
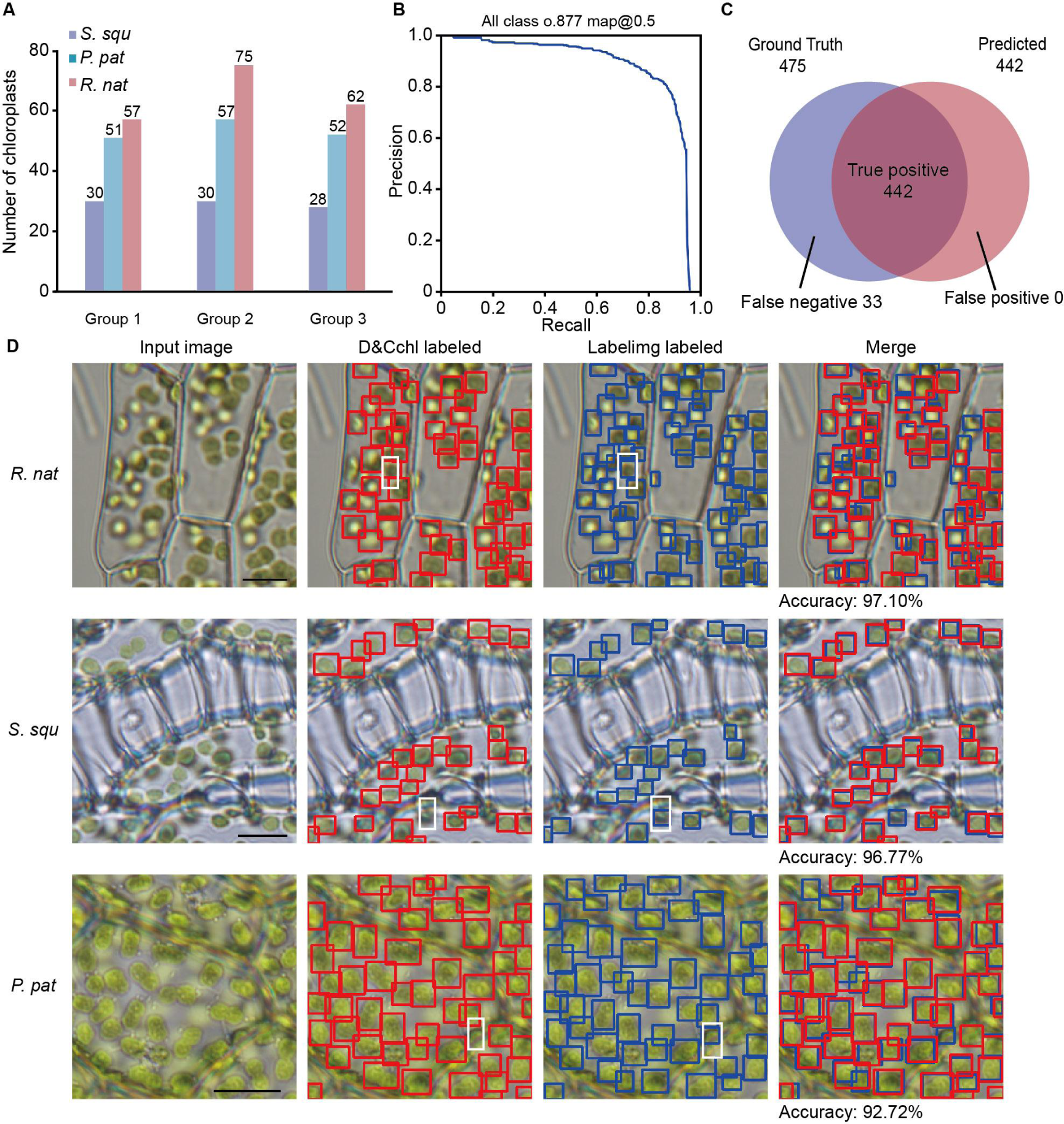
Application and accuracy assessment of the *YOLOv7-D&Cchl* model using 2D images. **A)** The results of applying the ***YOLOv7-D&Cchl*** model to count chloroplasts using 2D images from three plants, with three images for each plant. **B)** The precision-recall curve, effectively highlighting the model’s exemplary performance in both precision and recall. Notably, at a classification threshold of 0.5, the model achieves a precision of 87.7%. **C)** The Venn diagram visualization compares the ground truth, predicted, false negative, true positive and false positive detections of results from the ***YOLOv7-D&Cchl*** model, demonstrating the high accuracy and precision of the model. **D)** The detailed detection results of applying the model to detect chloroplasts using 2D images of three plants. Red boxes represent model detections, blue boxes represent manual detections, and the white boxes emphasize the missing detection by the model due to the off-focus of chloroplasts. Three plants include *Sphagnum squarrosum* (*S. squ*), *Physcomitrium patens* (*P. pat*), and *Ricciocarpos natans* (*R. nat*).

### Extended Usage of YOLOv7-D&C for Chloroplast Detection and Counting with 3D Images

We also noticed the mis-detected events and concluded that the white boxes in Figure 2B and C, due to chloroplasts at different positions not being simultaneously in focus, out-of-focus chloroplasts were not detected by the D&Cch model. To address this issue, a multilayer of light-microscope images covering a whole cell was collected (Fig. 1B; Fig. 3; Supplemental Fig. S8). To avoid duplicate counting, IOU calculations were performed between each detected target in the second image and the targets in the benchmark. The maximum value was selected and compared to the predetermined IOU threshold (set at 0.3). If the IOU exceeded the preset threshold, it was classified as an existing chloroplast in the benchmark to prevent duplicate counting, otherwise, it was identified as a new chloroplast and added to the benchmark (Fig. 3). Sequenced images of *S. squ*, *P. pat*, and *R.nat* leaf cells were obtained at various focal depths (Fig. 3A). YOLOv7-D&Cchl was applied for each individual layer, and IOU was used for monitor the overlaps of every target between layers. The accurate counts of chloroplasts in different plants were obtained, with the counts being 326, 184, and 365, respectively. The precision was significantly improved in multilayer statistics than that in the single-layer image, as only about 90% percentage of total chloroplasts could be detected in a single-layer image compared to total multilayer images. Usually, scientists performed chloroplast detecting using single-layer image, because it is exceedingly difficult and time-consuming to get rid of repeated chloroplast counting in multilayer images (Grishagin, 2015). We compared the efficiency of manual detection and counting for multilayer images with the YOLOv7-D&Cchl-mediated chloroplast detection and counting, and concluded that our strategy obviously saves more time than the manual method. Additionally, after resolving the out-of-focus issue, the cases of miss-detection by YOLOv7-D&Cchl were almost completely ignored. Thus, our 3D detecting and counting method not only addresses the counting errors caused by severe out-of-focus and defocusing issues in 2D counting but also significantly enhances efficiency, which is particularly meaningful for studying organelles within whole cells.

**Figure 3.**
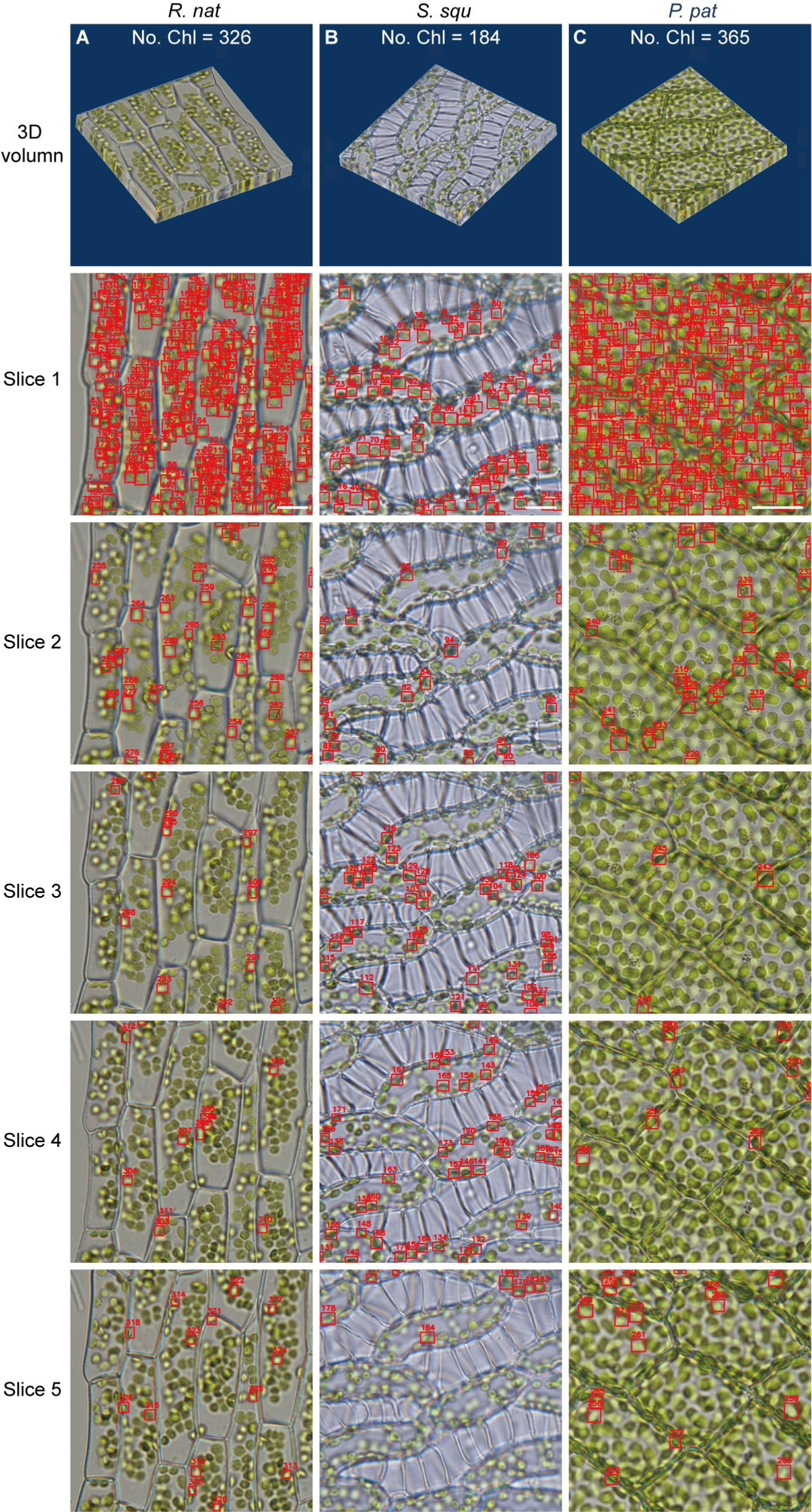
Application of the *YOLOv7-D&Cchl* model using series 2D images for 3D volume detection. **A)-C)** The microscopic image series and their chloroplast detection results for *Sphagnum squarrosum* (*S. squ*), *Physcomitrium patens* (*P. pat*), and *Ricciocarpos natans* (*R. nat*), respectively. Slice1-5 represented the first to the fifth layer of the image from each respective sample’s microscopic image series. Counting chloroplasts in a 3D volume was achieved by summing all the detected chloroplasts in these slices, while avoiding repeated counting. The detected chloroplasts were marked by red boxes. Each slice displayed only newly detected chloroplasts. Scale bars represent 10 μm.

### Association of YOLOv7-D&C with cell segmentation for chloroplast detecting in 3D single cell

Bio-scientists find single-cell counting to be more meaningful than counting based solely on single 2D images. Therefore, we employed a segmentation method, segmenting the data obtained from optical microscopy on a single-cell basis, and then carried out automatic segmentation. The pre-trained model ‘cyto2’ loaded into the segmentation tool *Cellpose* has been employed and slightly tuned for the cell segmentation. We re-trained the original model cyto2 with our dataset and obtained a new model named ‘cyto2pro’, and then utilized it to segment optical microscope images. We specifically chose intact individual cells as inputs for object detection, subsequently conducting chloroplast detection and counting within those individual cells (Fig.4). Single cells were segmented and numbered sequentially. As shown in Fig. 4B, chloroplasts were detected and counted in 4 cells. The numbers of chloroplasts detected in these cells were 24, 28, 37, and 25, respectively, for 2D detection (Fig. 4B; Supplemental Fig. S9). While conducting multi-layer counting at the 3D level, the results per cell were 35, 36, 51, and 32, respectively. Therefore, we can conclude that our 3D whole cell counting methods provide a more accurate reflection of chloroplast status in cells.

**Figure 4.**
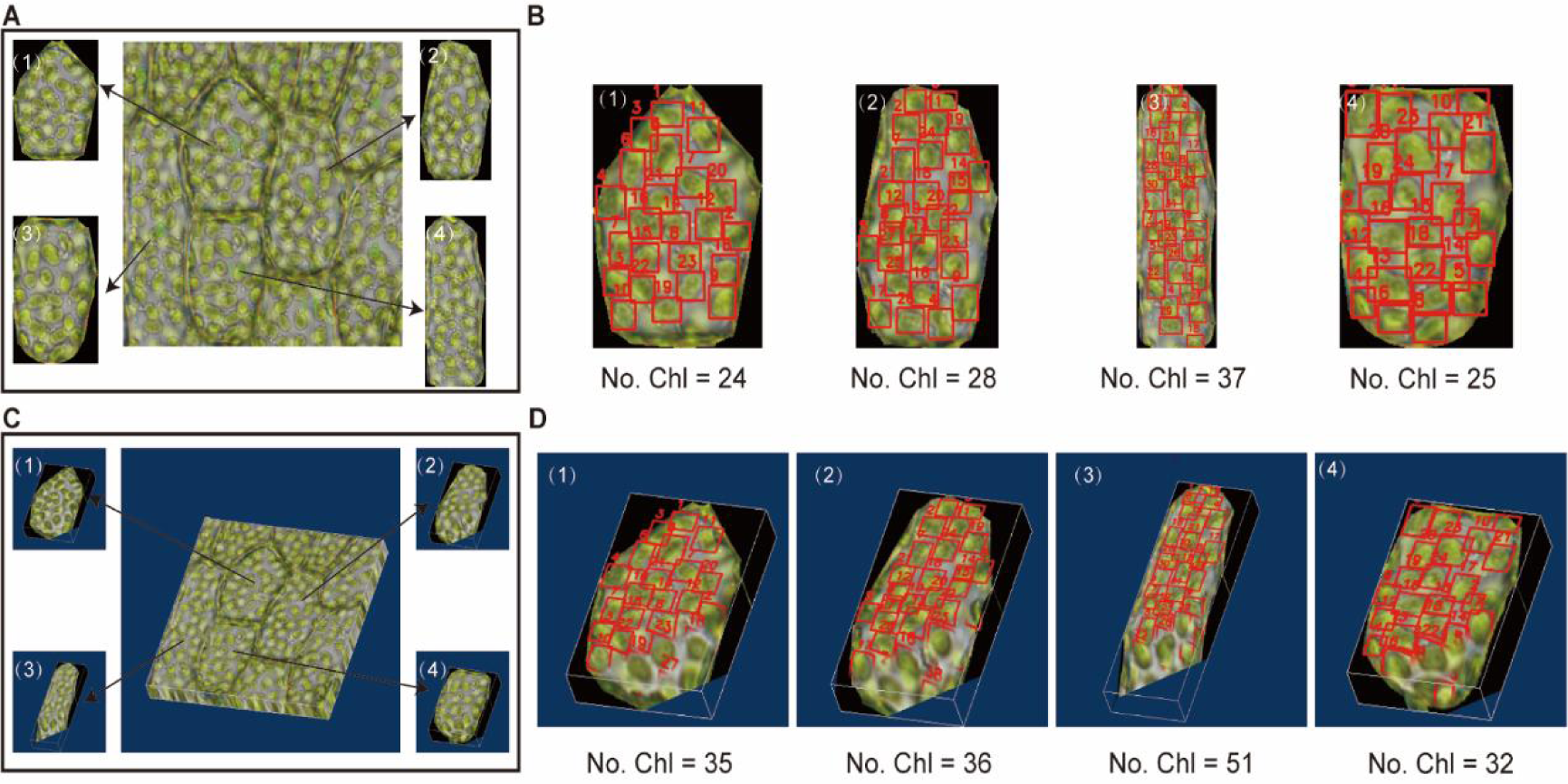
Chloroplasts Detection and Counting within 3D Single Cells. **A)** Single cell segmentation via ***Cellpose*** using single 2D images. The complete single cells were outlined for subsequent analysis. **B)** Chloroplasts detecting and counting via *YOLOv7-D&Cchl* in outlined 2D single cells. **C)** Single 3D cell segmentation via ***Cellpose*** using series 2D images spanning single cells, resulting a 3D single cell volume. **D)** Chloroplast detecting and counting via ***YOLOv7-D&Cchl*** in 3D single cells. For **B)** and **D)**, No.Chl showed the number of chloroplast detected. The red boxes showed the detected chloroplast.

Together, we developed the YOLOv7-D&Cchl as an excellent tool for chloroplasts counting. We also propose and utilize the IOU module to solve the duplication issue for counting multilayer-image stacks. Finally, with the integration of the cell segmentation tool Cellpose, single cell 3D counting has become a reality. This solution effectively addresses the challenges faced by our bio-scientists.

## Discussion

### The potential application of the D&C in plant cells

The innovative approach of D&Cchl has paved new avenues for plant cell research. Its potential applications extend to various fields in botany and cellular biology. Precise quantification of chloroplasts within plant cells can provide valuable insights into a plant’s photosynthetic capacity, growth, and developmental stages (Xiong et al., 2017; Fujiwara et al., 2019). By thoroughly understanding the quantity and distribution status of chloroplasts in different plant species or under varying environmental conditions, researchers can gain a deeper understanding of photosynthesis efficiency and the underlying factors behind it. Additionally, plants often undergo morphological changes during developmental process, as well as in response to environmental stresses like drought, extreme temperatures, or high light intensities, including adjustments in chloroplast distribution (Zahra et al., 2022; Liu X et al., 2018; Feng et al., 2019). Monitoring the number and distribution of chloroplasts in plant cells using D&Cchl benefits researchers by providing easy access to information regarding the plants’ status, growth rate, and developmental phase. Additionally, it helps in identifying environmental responses and understanding a plant’s adaptability to changing conditions.

### Our YOLO7-D&Cchl-IOU workflow for counting chloroplasts in 3D single cells is accurate and versatile

Currently, there are four main counting methods that are widely used, as depicted in Fig. 5A: manual counting, semi-automated counting, deep-learning-based DeepLearnMOR (Li et al., 2021), and the proposed D&Cchl. Manual counting is a straightforward method, but it is characterized by being time-consuming, labor-intensive, and susceptible to counting errors. For example, the ImageJ/Fiji provides several robust suite of counting tools, such as cell Counter (Plugins-Analyze-Cell Counter), to assist with manual labeling and auto-numbering (Arena et al., 2017). Semi-automated counting often requires user intervention and parameter configuration for image segmentation and object counting. For instance, the ImageJ/Fiji Analyze Particles (Analyze-Analyze Particles) allows one-click counting relying on image pre-treatment using threshold-based segmentation (Rueden et al., 2017; Schneider, et al., 2012), to ensure precision and dependability. However, it’s important to note that ImageJ has limitations in terms of counting accuracy, and it lacks batch processing capabilities. It can only process images one at a time. This single-image processing method may become time-consuming and less convenient when dealing with large-scale data analysis. DeeplearnMOR was developed based on YOLOv5 (YOLO version 5) for counting chloroplasts in fluorescent images. This approach offers the advantages of high precision and speed in two-dimensional object detection, as well as achieving full automation in counting (Tab. 1). However, the requirement of fluorescent images narrowed the practicability.

**Figure 5.**
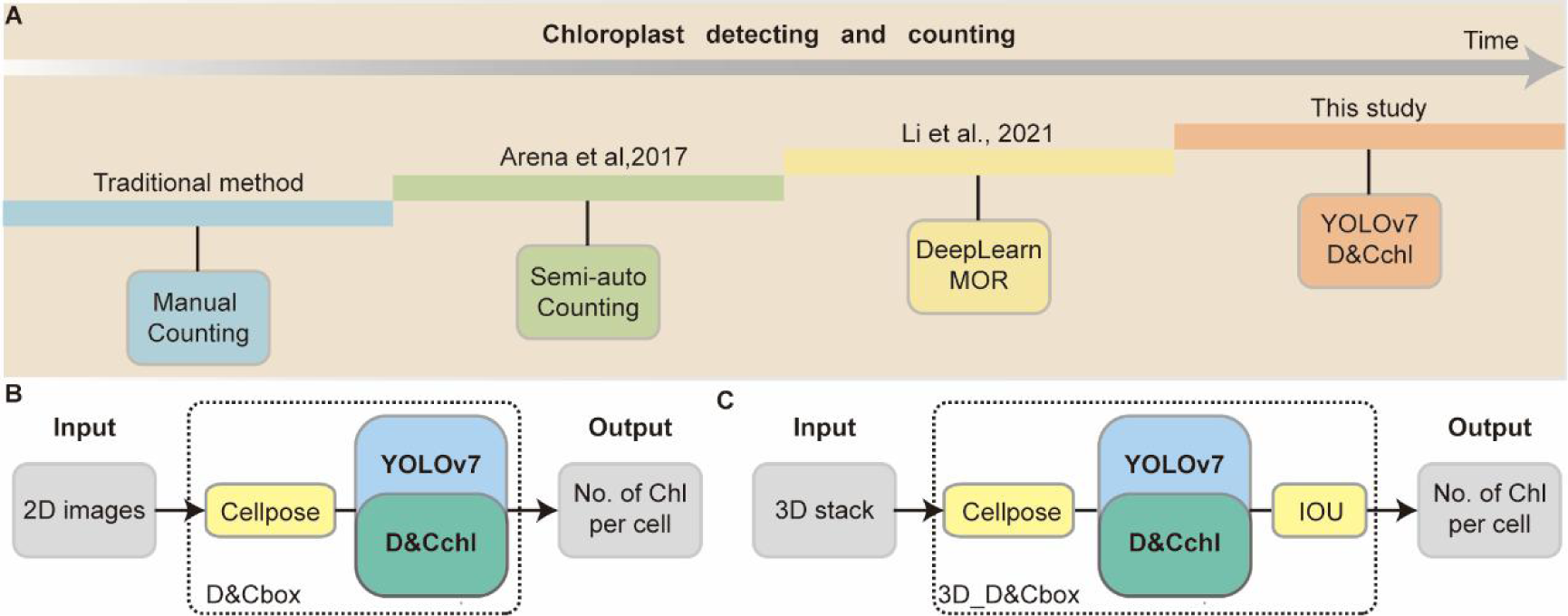
History of Chloroplast Counting Techniques Development. **A)** Overview of major methods for chloroplast counting, from manual to our deep learning-based 3D single-cell detection. **B)** Process of chloroplast detection in 2D cells. **C)** Workflow for chloroplast detection in 3D cells.

**Table 1.** Comparison of the four counting methods.

| Counting method | Scope of observation | Stained or not | Live Cell (Yes/No) | References |
| --- | --- | --- | --- | --- |
| Manual | 2D Slice | No | Yes | Lutz & Dzik,1993 |
| Semi-auto | 2D Slice | No | Yes | Arena et al., 2017 |
| DeepLearn MOR | 2D Slice | Yes | No | Li et al., 2021 |
| YOLOv7 D&Cchl | 3D, whole cell | No | Yes | This study |

Our proposed method for counting chloroplasts in 3D single cells not only achieves high accuracy, but also is versatile across various cell chloroplasts. As described in Fig. 5B and 5C, the 2D or 3D images captured via optical microscopes undergo **Cellpose** for cell segmentation and **YOLOv7-D&Cchl** for auto-detecting and counting (Fig. 5B, C; Tab.1). By utilizing this approach, we were able to achieve automated counting of chloroplasts, which has the potential for widespread use due to its straightforward data requirements.

### Extended usage of D&C associated with other deep learning-based tools

The D&Cchl method, rooted in the power of deep learning, offers a revolutionary solution for accurately counting chloroplasts in plant cells. Its value becomes even more significant when considering its potential synergy with other deep learning techniques. By integrating with object detection frameworks like Faster R-CNN (Faster Region-CNN) or SSD (Single Shot MultiBox Detector), D&Cchl can enhance the precision and segmentation accuracy of chloroplasts (Liu W et al.,2016; Girshick R et al.,2015). When paired with time series analysis tools like LSTM (Long Short-Term Memory), it becomes possible to monitor the dynamics of chloroplasts within plant cells across various environmental conditions in real-time (Hochreiter and Schmidhuber,1997). Furthermore, combining multimodal data such as spectral images and thermal imaging provides a comprehensive insight into the health status of plants. With the addition of technologies like autoencoders and VAEs (Variational Autoencoders), deep extraction of cellular features from microscopic images becomes feasible. In essence, the amalgamation of these technologies offers a more efficient and holistic approach to plant cell research.

In summary, different counting methods present their own merits and complexities. Our proposed approach overcomes the limitations of other techniques by harnessing the capabilities of deep learning and integrating various methodologies to achieve precise and comprehensive counting, offering a powerful tool for biological research.

## Methods

### Plant culture

Within a clean workbench, the plant’s surface soil was first rinsed off with clear water, followed by immersion in a 0.05% Triton buffer solution for 5 minutes. Subsequently, the specimen was treated with a 5% NaClO solution for another 5 minutes and then rinsed with sterilized distilled water three times, each rinse lasting 2 minutes. Excess moisture on the surface of the thallus was absorbed using sterile filter paper. The specimen was then placed on a pre-prepared ½ GB5 medium with 1% sucrose, sealed with a sealing film, and then incubated in a room maintained at a constant temperature of 22℃ with a light cycle of 16 hours (at 800 lux) to 8 hours. For cultivation, thallus sections that were vibrant green and in good growth condition were placed on the same medium for propagation. During this process, the growth status of the scales was observed and documented. The culture medium was prepared by dissolving Gamborg B5 Medium powder and sucrose in distilled water, adjusting the pH to 5.7-5.8, adding agar, and then autoclaving. After sterilization, the medium was poured into Petri dishes and allowed to air-dry at room temperature.

### Image collection

In this study, images of the samples were captured using an Olympus-BX43 biological microscope. Initially, we ensured the microscope slide was clean before adding 3-5 drops of distilled water onto it. Carefully, the liverwort scales were dissected using a dissecting needle and then laid flat in the water droplets on the slide. Once covered with a cover slip, we began our examination under the microscope at a low magnification to initially locate our target. Following this, we switched to a 10×60 high-power magnification for a more detailed observation. After confirming regions where the chloroplasts were relatively dispersed without excessive clustering, photographs were taken, thus yielding a high-quality dataset of chloroplast images, setting the foundation for subsequent analysis. By carefully adjusting the longitudinal translation stage of the sample stage, a series of the microscopic images focused at different depths of the cells were obtained to ensure that all the chloroplasts in the cells could be seen clearly and counted.

### Manual labeling to create the training dataset

In this study, we initially captured approximately 300 microscopic images using the Olympus-BX43 biological microscope. After selecting images that excelled in terms of resolution, contrast, and chloroplast distribution, we employed the annotation tool, labelimg, to manually label the chloroplasts present in these microscopic images. Utilizing the bounding box feature of this tool, we meticulously assigned a label to each visible chloroplast. In total, we labeled over 20,000 chloroplasts, of which about 90% were used as training data and the remaining 10% served as validation data. The annotated data is typically saved in txt format, providing precise training and validation datasets for subsequent deep learning training.

### Preprocessing and Training

To run YOLOv7 locally, the appropriate environment needs to be set up first. It’s recommended to begin by creating a dedicated Python environment using conda: conda create -n yolov7_env python=3.9, and then activate it with: conda activate yolov7_env. Within this environment, install the necessary libraries and dependencies using pip install -r requirements.txt. This encompasses the PyTorch deep learning framework and its vision libraries, as well as numerical computation and data processing libraries like numpy, scipy, and pandas, and data visualization libraries such as matplotlib and seaborn. OpenCV is also essential, being a computer vision library used for image processing and visual tasks. Once the dependencies are installed, clone the YOLOv7 project from its GitHub repository (https://github.com/WongKinYiu/yolov7). Proceed with further training or inference after obtaining the source code. Throughout the process, ensure all dependency libraries are correctly installed, and you may need to download pretrained model weights or other resources to run YOLOv7.

We trained the YOLOv7 network for 500 epochs using the Adam optimizer with a learning rate of 0.001 and a batch size of 16. The training process lasted 2.662 hours. All of our training and testing data were stored in an input directory which contained separate folders for training and testing, along with the corresponding label txt files. Our experiments were conducted on a desktop computer equipped with an Intel Core i7-10700 CPU @ 3.80 GHz and an NVIDIA GeForce RTX 3060 with 12GB of VRAM. The code was executed using the PyTorch 2.0.0 framework and was supported by CUDA version 12.2. Please refer to the supplemented manual for more details.

### Data analysis

We utilized manually labeled data as the ground truth reference. Any mislabeled or omitted annotations were meticulously counted by human inspection. The accuracy mentioned refers to the proportion of correct annotations made by the artificial intelligence system relative to the manual annotations.

The trained model can be evaluated using several metrics to measure its performance in the object detection task. These metrics include Precision (P), Recall (R), Average Precision (AP), and Mean Average Precision (mAP). The definitions of these metrics are as follows:

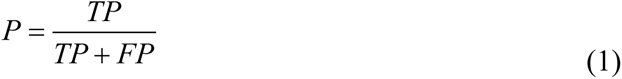

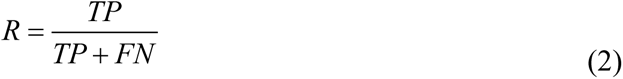

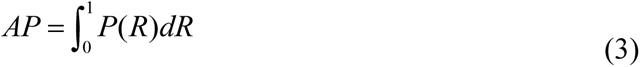

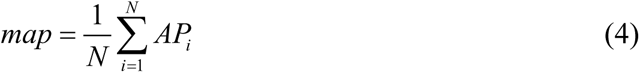

In this context, True Positive (TP) represents the correctly detected number of target chloroplasts in the image, False Positive (FP) indicates the count of misidentifications, and False Negative (FN) signifies the number of particles in the image missed by the network. Precision is a metric measuring the ratio of correctly detected particles out of all detected chloroplasts, reflecting the model’s accuracy in object detection. On the other hand, Recall measures the ratio of correctly detected chloroplasts out of all chloroplasts in the sample, showcasing the model’s capability in detecting all target instances. Average Precision (AP) is the mean precision value calculated across different recall levels, determined by the area under the precision-recall curve. AP offers insights into how well the model detects targets across various recall levels. Meanwhile, Mean Average Precision (mAP) is the average of AP values across all categories, providing an evaluation of the model’s overall performance.

### 3D model reconstruction

Our 3D image modeling was built upon a series of cell images captured at different focal depths under an optical microscope. To construct a comprehensive 3D cell model, we initially captured a sequential set of 2D images from the same sample, with each encompassing distinct depth information of the cell. These consecutive 2D image layers were then arranged in order of their focal depths, and sequentially stacked to form the cohesive 3D cell model. During this process, each 2D layer contributed structural information at a specific depth to the 3D model. To ensure the accuracy of the model, we employed the IOU technique to prevent chloroplasts from being counted multiple times during the stacking process. Through this approach, we successfully constructed a comprehensive 3D cell model from the series of 2D images, enabling more accurate chloroplast detection and counting in a 3D space.

### Code and software

All the code and training dataset have been shared on GitHub (https://github.com/xiaoli111111111/-AI4CELLBIO-ECNU), with detailed explanations in the supplementary manual.

## Supplemental information

Extended data (including 6 Supplementary Figures) is available for this paper.

**Supplemental Figure S1 The leaf of the three plant materials used for chloroplast detecting and counting.**

**Supplemental Figure S2 Network structure diagram of Yolov7 algorithm.**

**Supplemental Figure S3 Precision Curve**

**Supplemental Figure S4 Recall Curve**

**Supplemental Figure S5 F1 Score Curve**

**Supplemental Figure S6 Detailed Detection and Evaluation of Chloroplasts in Microscopic Plant Cells.**

**Supplemental Figure S7 Detailed Detection and Evaluation of Chloroplasts in Microscopic Plant Cells.**

**Supplemental Figure S8 Feasibility Analysis of 3D Chloroplast Detection.**

**Supplemental Figure S9 3D Single-Cell Detection Analysis.**

**Supplemental manual Codes used in the manuscript.**

## Funding Acknowledgement

We are thankful for the initial plant material provided by Prof. Rui-liang Zhu and Prof. Yue Sun. This research received support from the Natural Science Foundation of Hebei Province (No. F 2023402009), East China Normal University, and the National Natural Science Foundation of China (Grant No. 62175059).

## Author contributions

Z.D, Q.Z, N.J.L and S.Q. designed most of the experiments. S.Q performed most of the experiment. L.L, Q.Q.G and X.Y.L performed the plant material culture and collected image data. W.T, H.Z.S and W.H.Y assisted the workstation or platform preparation or others. Q.S, Z.D, N.J.L, S.K, and Q.Z wrote the manuscript and conceived the research.

## Data availability

The data underlying this article are available in the article and in its online supplementary material.

